# Using multiple reference genomes to identify and resolve annotation inconsistencies

**DOI:** 10.1101/651984

**Authors:** Patrick J. Monnahan, Jean-Michel Michno, Christine H. O’Connor, Alex B. Brohammer, Nathan M. Springer, Suzanne E. McGaugh, Candice N. Hirsch

## Abstract

**Background:** Advances in sequencing technologies have led to the release of reference genomes and annotations for multiple individuals within more well-studied systems. While each of these new genome assemblies shares significant portions of synteny between each other, the annotated structure of gene models within these regions can differ. Of particular concern are split-gene misannotations, in which a single gene is incorrectly annotated as two distinct genes or two genes are incorrectly annotated as a single gene. These misannotations can have major impacts on functional prediction, estimates of expression, and many downstream analyses.

**Results:** We developed a high-throughput method based on pairwise comparisons of annotations that detect potential split-gene misannotations and quantifies support for whether the genes should be merged into a single gene model. We demonstrate the utility of our method using gene annotations of three reference genomes from maize (B73, PH207, and W22), a difficult system from an annotation perspective due to the size and complexity of the genome. On average, we find several hundred of these potential split-gene misannotations in each pairwise comparison, corresponding to 3-5% of gene models across annotations. To determine which state (i.e. one gene or multiple genes) is biologically supported, we utilize RNAseq data from 10 tissues throughout development along with a novel metric and simulation framework. The methods we have developed require minimal human interaction and can be applied to future assemblies to aid in annotation efforts.

**Conclusions:** Split-gene misannotations occur at appreciable frequency in maize annotations. We have developed a method to easily identify and correct these misannotations. Importantly, this method is generic in that it can utilize any type of short-read expression data. Failure to account for split-gene misannotations has serious consequences for biological inference, particularly for expression-based analyses.

## Introduction

The annotation of a genome is a useful resource in many modern sequencing endeavors. It provides the initial link connecting mapping studies to functional impact, and it defines the context in which much of our genomic inference takes place. Modern software/pipelines (1) greatly facilitated production of *de novo* annotations for a large number of species, and multiple independent genome assemblies and annotations have been produced for more well-studied species (2–5).

Despite the importance of developing high quality annotations, and the exponential increase in annotated sequences over time that have come from assembly of many new genomes, the annotation process remains notoriously error-prone (1, 6, 7). Annotation pipelines attempt to integrate multiple data types, such as RNAseq, orthologous protein sequences, ESTs, as well as *ab initio* predictions from the genome sequence itself. In addition to the complexity of the data, the challenge is heightened by the complexity (and scale) of the underlying biological processes. Expression and maturation of transcripts and proteins is a highly dynamic process that varies over time as well as across different tissues, making it hard to differentiate between functional and intermediate forms. Furthermore, biological errors such as transcriptional read-through, as well as chimeric transcripts, provide conflicting evidence to the true underlying gene(s).

Research communities recognize the value of manual curation in the improvement of annotations and have encouraged input from community members (8, 9). Manual curation of gene annotations often comes from individual community members interested in a particular gene or gene family, relying on their detailed knowledge and data to identify and correct errors in a gene model. Depending on the community size and resource availability to a given study system, the extent to which this manual curation occurs and is effectively absorbed and corrected in future annotations is variable. Bioinformaticians can facilitate this process by developing automated algorithms that flag potential errors for subsequent manual curation.

The presence of multiple *de novo* genome assemblies and *de novo* annotations for a single species or multiple closely related species provides a useful dataset for such algorithms. By identifying the co-linear regions within each reference and linking the homologous genes across the annotations, researchers can discover discrepancies between gene models in the different genome assemblies. One particularly insidious discrepancy is when two distinct gene models in one annotation correspond to non-overlapping parts of a single, merged gene in the alternative annotation, commonly known as split-gene misannotation (10). These can have major impacts on functional predictions, estimates of expression, as well as downstream analyses. Here, we present a method to compare annotations and automatically detect potential split-gene misannotations, and subsequently determine which gene model (merged vs split) is likely correct, using transcript abundance estimates from short-read sequence data. Expression data from multiple tissues is standard input for most annotation pipelines (1), so in most cases, it should exist by virtue of having produced an annotation. This generic method accommodates all standard RNAseq libraries, including single-end and non-stranded preparations.

The difficulty of the annotation process, and thus the prevalence of errors, will vary greatly across study systems due to factors such as current and/or ancient polyploidy, transposable element (TE) content, and gene density throughout the genome. Maize is a good case system in which to test our misannotation detection method as it is an ancient polyploid with high TE content including TEs that are in close proximity to gene models. We analyzed *de novo* annotations from three maize genome assemblies, including W22 (11), B73 (12, 13), and PH207 (14). Using our pipeline, we identified hundreds of instances where multiple genes correspond to a single gene in an alternate annotation and determined the most likely annotation. We further demonstrate the biological misinterpretations that can result from these split-gene misannotations.

## Results

### Split-Gene Misannotation Detection and Classification Pipeline Overview

Our pipeline proceeds in two major steps: 1.) identification of potential split-gene misannotations (i.e. split-gene candidates) based on pairwise alignments (Figure 1; *Syntenic Homology Pipeline* in Methods) followed by 2.) determination of the supported gene model using short-read expression data (Figure 2; *Split-gene classification* in Methods). The output of the first step, which is based on a sequential alignment procedure using *nucmer* followed by reciprocal BLAST, is a key that labels the genes that have a one-to-one homologous relationship across the annotations along with the genes that have a one-to-many homologous relationship (a single gene in one annotation corresponds to multiple genes in the alternative annotation). The one-to-many genes will contain both tandem duplicates as well as split-merge candidates (Fig 1A). These two classes of one-to-many genes are distinguished by the proportionate overlap of the BLAST query genes with respect to the total aligned space of the subject gene (Fig 1B). The split-gene candidates are carried forward to the second “classification” step in the pipeline.

**Figure 1.**
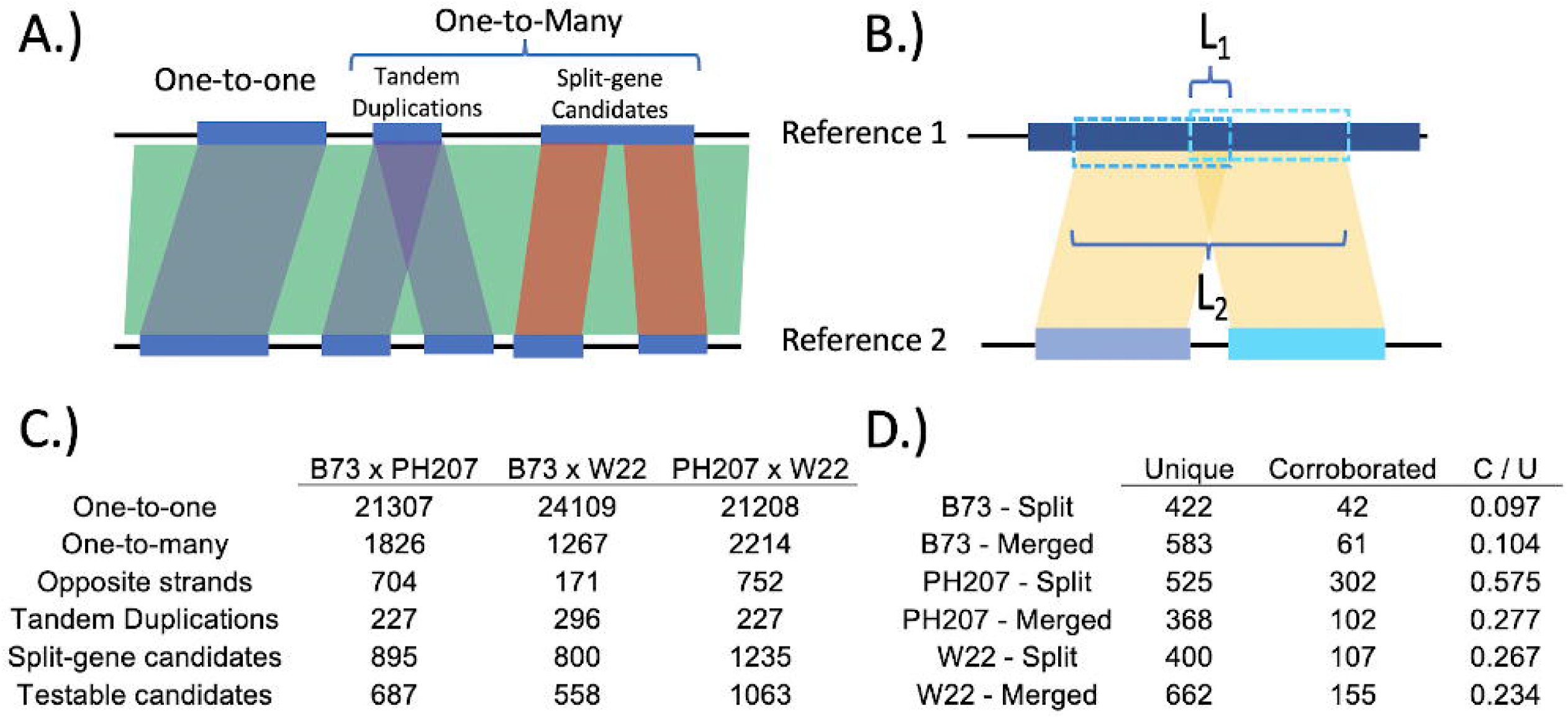
Identifying syntenic homologs and isolating split-gene candidates. A.) Homology classifications from syntenic homology pipeline. B.) Schematic for calculation of tandem duplicate percentage. We require the ratio of L1 to L2 to be < 0.1 (i.e. the proportionate overlap of the BLAST query genes with respect to the total aligned space of the subject gene.) C.) Summary of homology classifications and split-gene candidate filtration. A ‘Testable candidate’ is one in which all of the genes involved are expressed. D.) Corroboration of testable candidates. E.g. 43 ‘Corroborated’ split-gene candidates in the B73 annotation (“B73 - Split”) were simultaneously identified as a single gene in W22 and PH207, while there were 61 genes in B73 that corresponded to multiple genes in both PH207 and W22 (“B73 - Merged”), and the 438 ‘Unique’ split-gene candidates in B73 were identified as a single gene in W22 or PH207.

**Figure 2.**
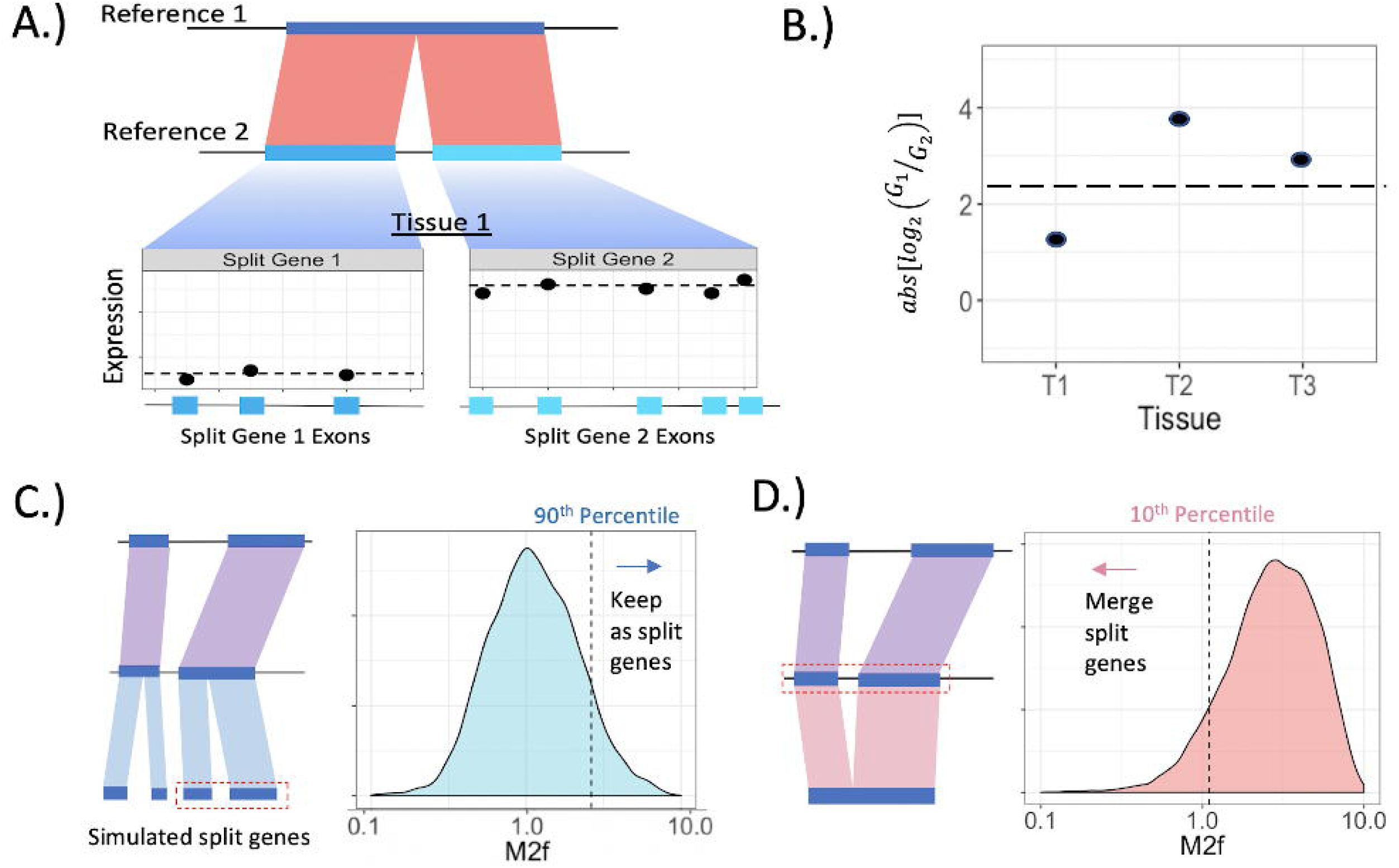
M2f approach for determining correct gene model(s) for split-gene candidates. A.) Calculating average normalized expression across exons within a tissue for a pair of split-gene genes. B.) M2f calculation. The absolute log_2_-fold change in average expression (from A.) across the split-genes is averaged across tissues. Higher values reflect large expression differences across split-genes. C.) Simulating the M2f distribution under the null hypothesis that split-gene expression differences come from a single underlying gene. Observed M2f values greater than the 90th percentile of this null distribution are unlikely to result if the single gene annotation is correct. D.) Simulating the M2f distribution under the null hypothesis that split-gene expression differences come from separate, adjacent genes.

Our classification method is based on the expectation that the difference in expression across the split genes should be greater if split (multiple) gene annotation is correct than if the merged (single) gene annotation is correct. To evaluate this degree of difference in expression patterns across the split genes, we developed the M2f (“Mean 2-fold split-gene expression difference”) metric (Fig 2A-B). Simulated, empirical null distributions (Fig 2C-D) are then used to determine significance thresholds for the M2f metric, based on if the value is lesser or greater than expected by chance. In other words, are the expression differences across the split-genes consistent with an underlying biological reality of a single gene or multiple, distinct genes?

To demonstrate the utility of this identifcation and classification method, we analyzed three maize reference genome assemblies that each of been independently annotated. The annotations under consideration represent different stages of development as well as different types and amounts of validating data. The annotation for B73 is currently in its fourth version, whereas W22 and PH207 are in their second and first version, respectively. Annotation of B73 was based on six evidence types, including long-(PacBio IsoSeq) and short-read RNAseq, optical mapping, full length cDNAs (from BACs), and orthologous protein sequences (12). The IsoSeq expression data from B73 was also utilized for annotation of W22 as well as short read data and optical mapping specific to W22 (11). The PH207 annotation only included standard short-read RNAseq data from PH207 (14). All annotations were produced using the MAKER-P pipeline (15) (with a modification for long-read expression data for B73 and W22) and contain approximately the same number of genes (~40k). Due to the lesser data used for the genome and annotation of PH207, the completeness and accuracy are predictably lower for PH207.

### Identification of Maize Candidate Genes

Alignments generated using *nucmer* covered a large portion of the genome with the greatest total alignment length between B73 and W22 (1.07 Gb; ~46%). Pairwise alignments with PH207 covered a much lower (~37%) proportion of the genome, regardless of whether it was aligned to B73 or W22. Furthermore, the alignments with PH207 were broken up into many smaller aligned regions (~60% of the average length in B73 x W22; Supp. Table 1). From the syntenic homology pipeline (Figure 1A) for each pairwise comparison, we found >20k one-to-one homologs (with the greatest number identified in the B73 x W22 comparison, likely due to the shared IsoSeq data) and 1.2 - 2.3 thousand instances of one-to-many homology across the pairwise comparisons (with the greatest numbers identified for comparisons involving PH207; Fig. 1C; list of one-to-one and one-to-many homologous genes in Supp. File 1 and 2, respectively). The dominant scenario was for multiple genes in PH207 to correspond to a single gene in either B73 or W22. However, in 37% (comparison to B73) and 44% (comparison to W22) of these instances, the split PH207 genes were on opposite strands, and and often overlapping (Supp. Table 2), perhaps indicative of overannotation of antisense transcription events in PH207. Such opposite and overlapping split-genes were also observed in B73 and W22, but to a much lesser extent (Supp. Table 2).

Nonetheless, after filtering the remaining candidates to remove possible tandem duplications and retain only expressed genes, there remained substantially more split-gene candidates (Corroborated + Unique = 507 + 307 = 814; Fig. 1D) in PH207 versus B73 (481) and W22 (525). Furthermore, the number of split-gene candidates in PH207 that were found to correspond to a single gene in both B73 and W22 (i.e. they were ‘Corroborated’; Fig 1D) is much higher than the ‘Corroborated’ B73 and W22 split-gene candidates combined. This is again concordant with comparatively less data used for the PH207 annotation, where for example, a lowly-expressed gene in PH207 might lack the coverage necessary to generate a full-length assembled transcript, resulting in annotation of multiple genes instead of the single, true gene.

Considering these split-genes along with the merged genes to which they correspond, our analysis concerns 1275, 1383, and 2125 genes in the W22, B73, and PH207 annotations, respectively, corresponding to roughly 3 - 5% of all genes contained in these annotations. Attributes of these genes tend to be comparable in many regards to the one-to-one homologous genes, except that they are usually nearer to neighboring genes and show more tissue specific expression (Supp. Fig 1).

### Classification of Maize Split-Merge Candicate Genes Using the M2f Metric

For each of the split-gene candidates identified with the syntenic homology pipeline (Fig1A), we sought to determine the gene model(s) with greatest support (i.e., should the split-genes remain split or be merged into a single gene?) using our M2f metric. The observed distributions of M2f for the split-gene candidates from each annotation are presented in Figure 3A, along with the threshold values (dotted lines) from the simulated, null distributions. We observe clear differences in the overall distributions of the M2f metric across the different genotypes (Figure 3A, Table 1), which leads predictably to differences in the number of split-gene candidates classified as either *merged* (i.e., the annotation in which the split-genes were annotated as a single gene is supported) or *split* (i.e., the separate, split-gene annotation is supported) (Figure 3A-B). The list of split-gene candidates, along with the supported annotation, are provided in the Supp. File 9.

**Figure 3:**
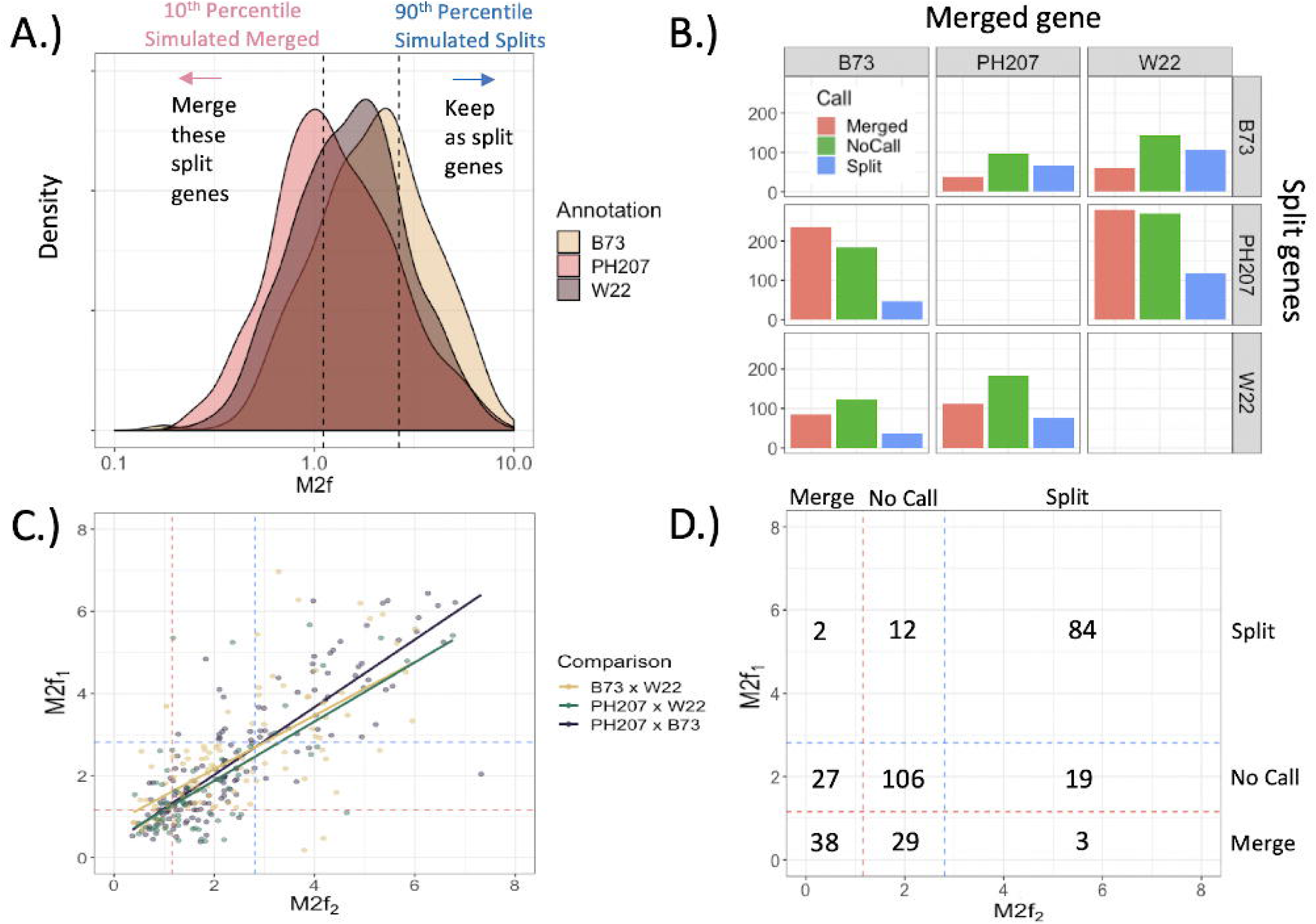
Results of M2f classification. A.) Observed M2f distribution across all split-genes detected in each annotation. The dotted lines are the threshold values generated by simulating null distributions in Figure 2 (C - D). B.) Number of split-gene candidates (Multiple genes) classified as to whether the split-genes should be annotated as distinct genes or a single, merged gene for each pairwise comparison of annotations. C.) Correlation of M2f values for instances where a single gene from one annotation corresponded to split-gene candidates in both of the alternative annotations (‘Corroborated’ Merged genes in Fig. 1D). E.g. Each point in the “B73 x W22” comparison corresponds to a single PH207 gene. X-axis is the M2f value from the B73 split-gene candidate, and y-axis is the M2f value from the W22 split-gene candidate. Dotted lines indicate the M2f threshold values in part A. D.) Joint distribution of classifications across comparisons in part C.

**Table 1.** Summary of M2f distributions for split-gene candidates in each annotation. CV=coefficient of variation. N=number of tested candidates.

| Split-genes | Mean | Median | Variance | CV | N |
| --- | --- | --- | --- | --- | --- |
| B73 | 2.45 | 2.09 | 2.49 | 0.693 | 506 |
| PH207 | 1.64 | 1.2 | 2.07 | 0.88 | 1129 |
| W22 | 2.05 | 1.66 | 2.42 | 0.759 | 614 |

The M2f distribution of split-gene candidates in the PH207 annotation (the lowest quality annotation, which make up a majority of the overall split-gene candidates) is shifted left relative to the other annotations (Figure 3A, Table 1), indicating that many of these are likely misannotations and should be merged as they have been annotated in either W22 and/or B73 (Figure 3B). Out of the 1129 sets of split-gene candidates in the PH207 annotation that were identified in either the comparison with B73 or W22, we found 505 that should be merged versus only 162 that should remain as separate genes. We were unable to make classification for 462 candidate sets based on the 10th and 90th percentiles of the simulated distributions. We observed the opposite pattern for split-gene candidates in the high-evidence B73 annotation (96 split-genes should be merged, 170 should remain as separate genes despite being merged in PH207 or W22, and 240 were unable to be called), where the separate gene models tended to have higher support based on M2f. The B73 gene model(s) tended to be favored by the M2f metric overall in comparison with either W22 or PH207, in line with B73 having the deepest evidence sources used to develop the annotation.

Having multiple pairwise comparisons also allows us to determine the consistency of the M2f metric. We consider instances where a single gene in one annotation corresponded to multiple genes in both of the alternative annotations. This provides two M2f values for this single gene, which should be correlated if M2f is sensitive to the underlying biological truth. In Figure 3C, we plot this correlation in M2f metrics for each annotation. In this plot, the “B73 x W22” correlation concerns the single PH207 genes that correspond to multiple genes in both B73 and W22. We find this correlation is highest when W22 is the annotation with a single gene corresponding to multiple genes in both PH207 and B73 (B73 vs. PH207 correlation = 0.85), followed by B73 (PH207 vs. W22 correlation = 0.68) and PH207 (B73 vs. W22 correlation = 0.66). While these correlations are imperfect, they rarely lead to conflicting classifications (Figure 3D) and, typically, the M2f value trends in the same direction even if the gene model does not pass the null distribution thresholds. Of the 320 instances where a single gene corresponds to two or more split-genes in both of the alternate annotations, only five (1.56%) are in conflict (i.e. M2f supports merging the split-genes for one of the alternative annotations, while the other alternative annotation suggests the genes should be kept separate, or vice versa; Figure 3D).

To further test the robustness and validity of our approach we investigated a number of potential confounding factors (Supp. Figs 2-4) that could impact classification of genes based on the M2f metric. First, we examined if genes that produce multiple isoforms have inflated M2f values. We compared the M2f distributions for B73 genes with multiple isoforms versus single isoforms (Supp. Fig 2) and found a slight inflation of M2f values for genes with multiple isoforms (Median M2f of 1.41 vs 1.59 for single and multi-isoform genes, respectively, within the split-gene candidates). Although this bias is slight, it serves to emphasize the role of the simulations in protecting against potential artifacts. As long as the simulated data is representative of our split-gene candidates (multiple isoform genes, in this case, are not over-represented in our candidates), the simulated null distribution will include this M2f inflation, thus protecting against misclassification due to this artifact. Notably, in our study, multi-isoform genes within our B73 candidates are less frequent in the empirical data (0.54) than to either the simulated split genes (0.64) or the simulated merged genes (0.59). We also explored the impact of exon number on our M2f metric and found that there is little impact of exon number on the distribution of M2f values (Supp. Fig 3). Finally, we explored the impact of using annotations from the different genome assemblies to set the thresholds for setting the 10^th^ and 90^th^ percentiles, and found the thresholds were highly similar across the genomes (Supp. Fig 4).

### Features of Classified Maize Genes

We explored features of the classified genes to determine if there were common features that could be informative in improving future automated annotation efforts. Genes that were originally annotated as a single/merged gene model but were determined to be split based on the M2f metric tended to be longer (Figure 4B) and have more exons (Supp. Figure 6A). Merged gene models supported by our M2f metric (MS = merged supported) were longer than the misannotated, merged genes (MNS = merged not supported); yet, MS genes have comparatively fewer exons than MNS genes (Supp. Fig 6A and C). The long, exon-sparse MS genes may be more likely to be missing reads spanning particular exon-exon junctions and, thus, be more prone to being misannotated as multiple genes (particularly when relying on short-read RNAseq data).

**Figure 4.**
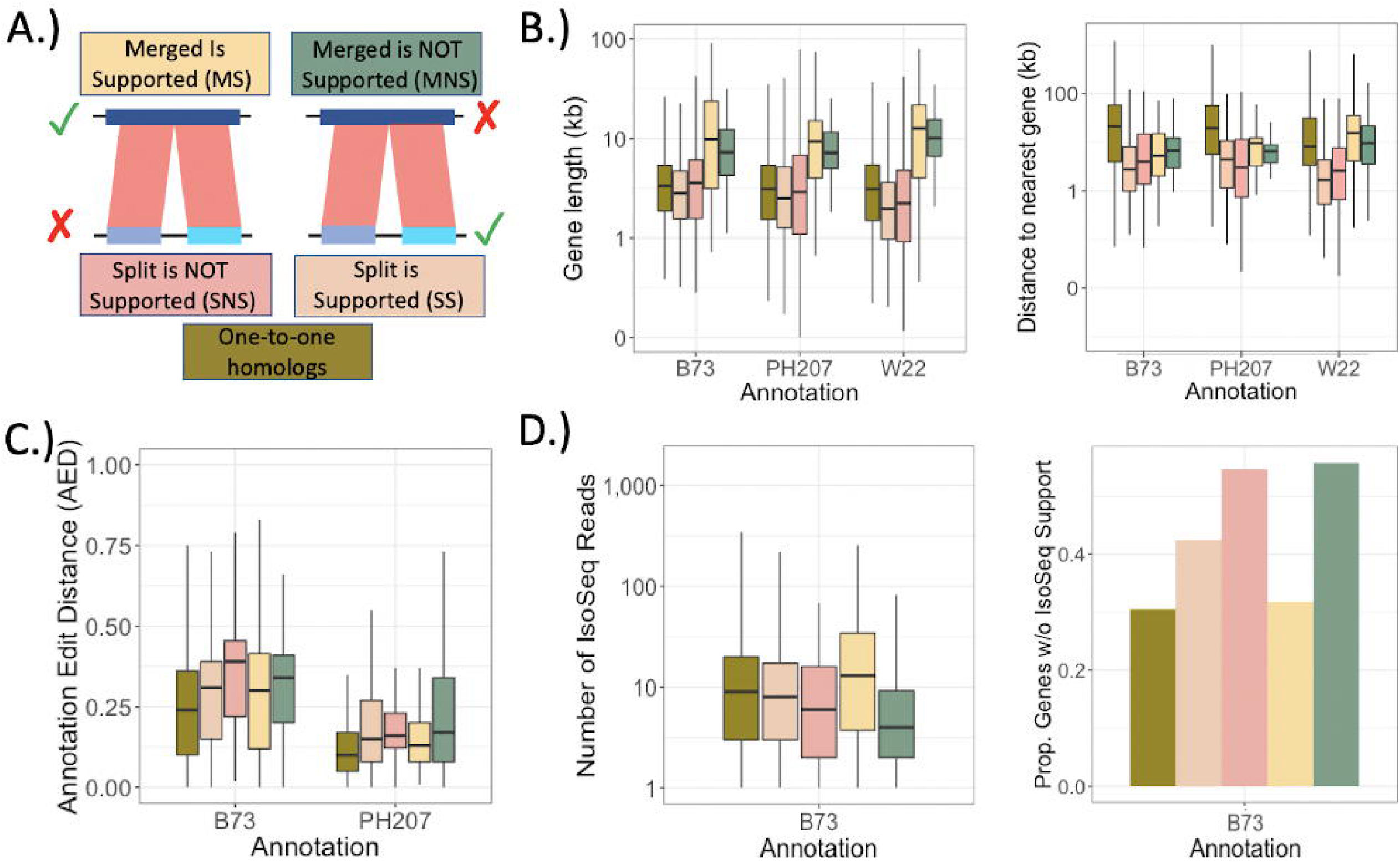
Features of one-to-one genes as well as split-gene candidates. A.) Split-gene candidates are classified based on whether they were initially annotated as split or merged for a given genotype followed by the classification based on the M2f method. E.g. The “SS” box for the B73 genotype are instances where multiple genes in B73 correspond to a single gene in either PH207 or W22, and the multiple (split) genes of B73 were determined to be the correct annotation. Outliers were removed on all plots. B.) Length and Distance between genes. C.) AED calculated from MAKER-P for the B73 and PH207 annotations. For B73, multiple isoforms were annotated, and we took the max AED across all isoforms for a given gene model. D.) Number of IsoSeq cDNAss for genes in each category. Genes with no IsoSeq support were excluded and shown separately as a proportion on the right. IsoSeq cDNAs were filtered for MQ > 20 and for coverage of at least 75% of the longest transcript sequence.

Generally, the split-gene candidates (including genes originally annotated as split, along with their merged counterparts in the alternate annotations), tend to be closer to other genes as compared to the genes with one-to-one homology across all three annotations (median distance of 3.6kb versus 4.1kb). This suggests that gene dense regions may be more prone to split-gene misannotations, and that these misannotations may be more frequent in species with smaller, gene-dense genomes. Looking within the split-gene candidates (all categories except for “One-to-one” in Figure 4), we found that when split gene annotation is supported, the components of the unsupported merged gene tend to be closer together. This suggests that the distance between these components contributed to the misannotation as a merged gene, potentially through a mechanisms like transcriptional read through of proximate genes. We observed the opposite trend in the PH207 annotation, but only for the split-genes in PH207 that correspond to a single gene in W22 (SNS distance = 3.6kb; SS distance = 5.3kb).

We also investigated whether expression differed between supported and unsupported annotations. Overall, expression abundance did not markedly differ from that seen in the one-to-one genes (Supp. Fig 6A). One slight exception is for the genes that were incorrectly annotated as a single, merged gene (MNS), where there is a higher density of high expression for these “genes.” Increased expression of one or multiple proximate, distinct genes may increase the likelihood of producing chimeric transcripts (e.g. via transcriptional read through), thus promoting incorrect annotation as a single, merged gene. Tissue-specificity of expression differed markedly between classification categories (Supp Fig 5B, D), particularly for the highly tissue-specific genes (Sup Fig 5D). We found that split-gene annotations (both SS and SNS) were more likely to result when expression of one of the genes was highly tissue-specific, whereas merged gene annotations (both MS and MNS) occured more often when expression was less tissue-specific. Interestingly, within each of these categories, the subset of supported annotations (as determined by our M2f metric) tended to be more tissue-specific than the nonsupported annotations (Supp. Fig 5D).

The Annotation Edit Distance (AED) is a common annotation quality metric that can be used to summarize the differences between an annotated gene model and the supporting evidence (16). We find that the AED reported by MAKER-P for the B73 and PH207 annotation is consistently higher for split-gene candidates as compared to the one-to-one homologs (Fig 4C), indicating lower quality of these gene models, generally. Notably, the AED of nonsupported annotations (SNS and MNS) is higher than the supported annotations (SS and MS). However, the AED distributions of supported and nonsupported split-gene annotations are largely overlapping; thus, while AED is sensitive to split-gene misannotation, it cannot be used to robustly identify incorrectly merged or split gene models.

We find that nonsupported annotations in B73 have lower or no IsoSeq coverage as compared to supported annotated gene models (Fig 4D). Both of the nonsupported annotation categories (SNS and MNS) have the highest proportion of genes with no long-read support (SNS = 0.54 and MNS = 0.58 versus SS = 0.42 and MS = 0.32). When we consider only the genes that have long-read support, there tend to be fewer supporting reads for the nonsupported annotation categories, particularly when B73 has an nonsupported, merged gene that M2f suggests should be split (Median number of IsoSeq cDNAs for MNS = 4 and SNS = 7 versus MS = 11 and SS = 8).

### Consequences of Split-Gene Misannotations on Biological Findings

We explored the consequences of split-gene misannotations for biological inference that rely heavily on the annotation, namely expression-based analyses. Comparing across genotypes, we found that genes that are one-to-one homologs show a much tighter correlation in normalized expression (r = 0.92) than the correlation between supported split-genes and their corresponding (nonsupported) single, merged gene(r = 0.43; Figure 5A; SS category in Figure 4). If two distinct genes are incorrectly annotated as a single gene, the estimated expression for the single gene will be an average of the expression of the two loci. Unless the two loci happen to be expressed similarly, this average will likely be more dissimilar from either of the two distinct genes than if we were to compare expression with the true homologs (i.e. if the misannotated merged gene was correctly annotated as two distinct genes). The dissimilarity may be further amplified by normalization procedures that scale read counts by the length of the feature over which expression is being measured. For an equivalent number of reads, the longer, merged gene model will have lower normalized expression. On the other hand, when the single, merged gene was supported, we found a very tight correlation between the expression of this gene and the corresponding (nonsupported) split-genes (r = 0.99; Supp. Figure 7).

**Figure 5.**
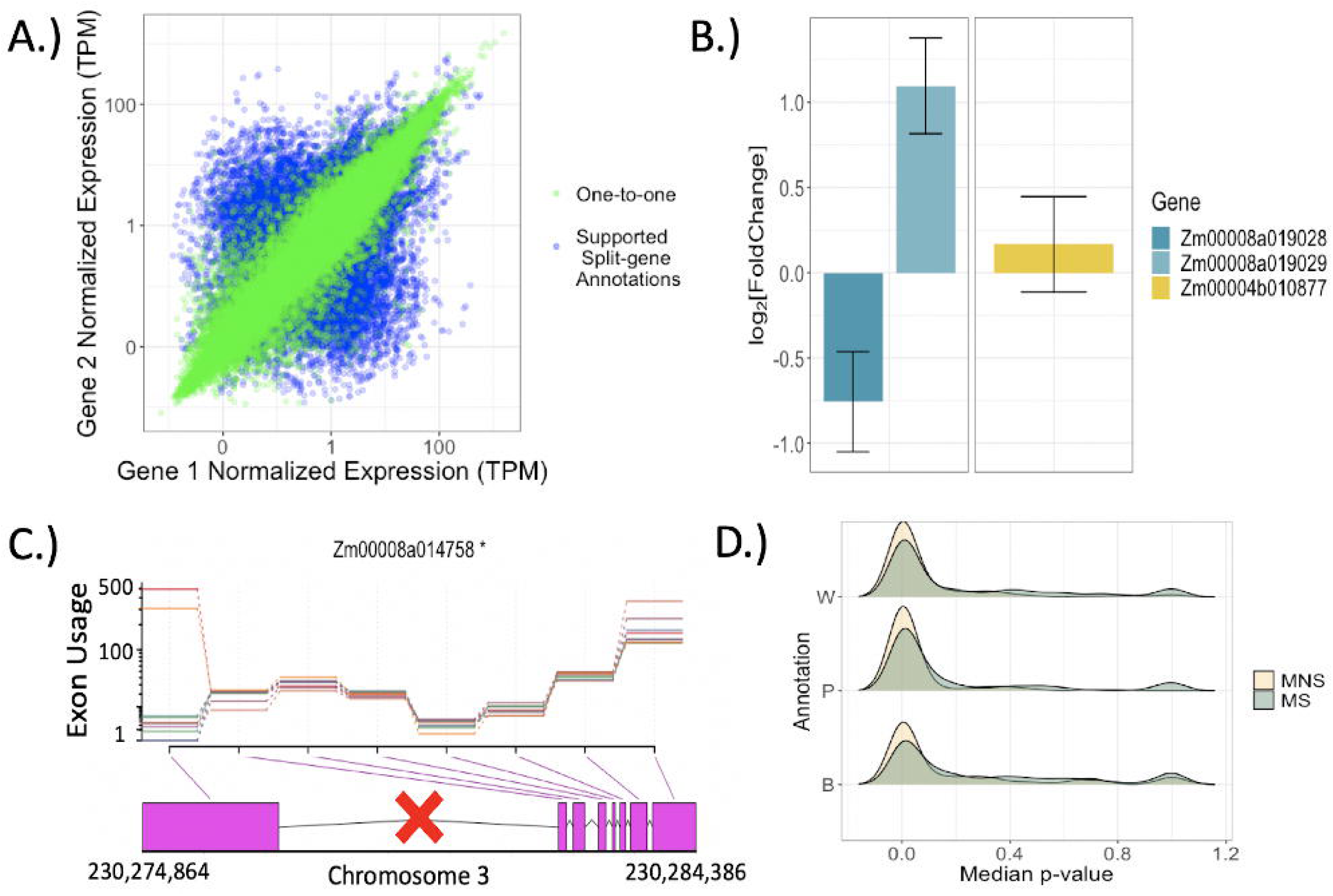
Consequences of split-gene misannotations. A.) Comparing expression estimates across homologs. For the correct split-gene annotations (Split Supported (SS) in Figure 4), expression of each split-gene is compared to the one expression value from the single gene that they correspond to. B.) Exemplar of differential expression misinference when two distinct genes are incorrectly annotated as one. Expression differences (between immature ear and anther) of component genes cancel out resulting in no differential expression for the single rightmost gene. C.) Example of misinference for differential exon usage. Incorrect annotation as a single gene in PH207 should be two genes (split at location demarcated with the red X) as annotated in W22. Colored lines indicate separate tissues. D.) Median p-value across the per-exon tests of differential exon usage for each gene. Inflation of low p-values is observed when distinct genes are incorrectly treated as a single, merged gene (Merged is not supported (MNS) in Figure 4).

Poor estimations of transcript abundance for split-gene candidates presumably will have consequences on inference of differential expression as well as differential exon usage. For example, the two PH207 genes in Figure 5B are differentially expressed albeit in opposite directions across the immature ear and anthers, yet these differences cancel out when we test for differential expression of the single, merged gene as annotated in W22 (Figure 5B). Similarly, Figure 5C illustrates improper inference of differential exon usage of the left-most exon in two of the tissues, when in fact, this exon is a distinct (and differentially expressed) single-exon gene according to our results. Across all of the nonsupported merged genes, there is an abundance of differential exon usage as compared to the supported merged genes (Figure 5D), suggesting that unsupported merged gene models lead to false inference of differential exon usage. We also observed this trend for the DEseq analysis, albeit to a lesser degree (Supp. Figure 8). A much higher proportion of exons are inferred to be differentially used across tissues for these nonsupported gene models, which is expected when the nonsupported merged gene is composed of two or more multi-exon genes (Supp. Figure 9). Therefore, these types of misannotations are highly predisposed for misinference of underlying biological processes.

## Discussion

Accurate gene models are of paramount importance in the era of genomics. While the bioinformatics community continues to develop and improve tools for the prediction of gene models (i.e. annotation), the burden of verifying and, if necessary, correcting these predictions is largely spread across the individuals invested in researching the particular organism. Bioinformaticians can do more to facilitate this process by developing methods that flag/correct misannotated genes, preferably without requiring the generation of additional data. We have described a comparative approach to identify potential split-gene misannotations across annotations of individuals within a species or closely related species, and a method to infer the correct annotation using pre-existing RNAseq data.

Though our approach is based on short-read RNAseq data, the utility of long-read expression data is clear in our results. PH207, which was the only annotation that did not utilize long-read data, exhibited substantially more split-gene misannotations than W22 and B73 combined. A single long read can capture all of the exon-exon junctions, whereas observations on one or more junctions are more likely to be missing with short-read sequencing due to random variation in sequencing coverage. In line with this, we find split-gene misannotations are more often associated with lowly-expressed and/or tissue-specific genes.

We have, however, shown that even annotations that are based on long-read data will still contain split-gene misannotations. These misannotations are not due to the long-reads *per se*; they more likely result whenever long-read data is unavailable for a particular gene and short-reads are sparse (e.g. lowly expressed genes). Our method capitalizes on the fact that these same genes may be more highly expressed in other genotypes, thus providing more complete evidence of the underlying gene model. The more salient issue with long-reads (and short-reads, for that matter) is the potential for aberrant transcriptional readthrough events that encourage improper merging of adjacent gene models (17). Fortunately, such events ought to be detected by our method, as these merged genes will more likely show highly inconsistent expression patterns.

In its current implementation, our method will not detect all instances of split-gene misannotations. Thus, we may underestimate the abundance of split-gene misannotations. The most obvious cause of non-detection would be if the gene(s) were consistently misannotated across all of the annotations being compared, in which case we would identify these genes as one-to-one homologs. However, by increasing the number of independent annotations considered, we should increase the odds that at least one annotation possessed the correct gene model. We also are only considering split-gene candidates where both of the split genes are expressed. Our attempts to handle the 0-expression genes introduced clear artifacts in our M2f metric, though an alternative or modified metric could possibly accommodate these scenarios. Lastly, we cannot strictly discriminate between truly split genes and certain scenarios of a single gene with multiple isoforms. Our simulation framework will partly protect against M2f inflation from multiple isoforms, since multiple isoform genes were well-represented in the simulated split or merged genes. However, a multi-isoform gene in which the predominant isoform is simply a truncated version of the longest isoform may still result in false positives via our approach. For these reasons, we view our method as a high-throughput means of flagging potential misannotations and suggesting the correct gene model, in order to facilitate the manual curation process of the larger community (8, 9).

## Conclusions

In summary, as additional *de novo* genome assemblies and annotations are produced, the greater the opportunity to identify and correct errors and inconsistencies. We have described a method to facilitate this process for split-gene misannotations, which we have demonstrated can strongly bias a range of biological estimates. Given that the required input (RNAseq) is readily available by virtue of having produced an annotation, our method could be integrated as a standard part of the annotation process for systems in which annotations already exist for other genotypes or individuals. Accrual of such tools are an important step towards developing accurate and consistent genome annotations, a foundational resource in the age of genomics.

## Methods

### Maize Datasets

We focus on three maize genome assemblies and corresponding annotations for this study: B73 (version 4; AGPv4, ftp://ftp.ncbi.nlm.nih.gov/genomes/genbank/plant/Zea_mays/latest_assembly_versions/GCA_000005005.6_B73_RefGen_v4) (12, 13), W22 (version 2, ftp://ftp.ncbi.nlm.nih.gov/genomes/genbank/plant/Zea_mays/latest_assembly_versions/GCA_001644905.2_Zm-W22-REFERENCE-NRGENE-2.0) (11), and PH207 (version 1, http://dx.doi.org/10.5061/dryad.8vj84) (14). We filtered annotations for a single gene with multiple transcripts by filtering for the longest coding sequence (CDS). We then converted these representative transcripts to fasta format and created a BLAST database (18) for each reference.

For each of the three genotypes, we collected tissue from ten spatiotemporally diverse tissues including: primary root six days-after-planting (DAP; R), shoot and coleoptile six DAP (SC), internode at vegetative stage 11 (V11; I), middle of the 10^th^ leaf (V11; L10), leaf 30 DAP (L), meiotic tassel at vegetative 18 (V18; T), immature ear at V18 (IE), anthers at reproductive stage 1 (R1; A), endosperm 16 DAP (En), and embryo 16 DAP (Em). These tissues were chosen to broadly capture variation in the maize transcriptome based on the maize B73 gene atlas (19). We collected two biological replicates per genotype/tissue combination and standard, non-stranded RNAseq libraries were prepared for each tissue sample using the Illumina TruSeq library preparation protocol-replicate.

Libraries were sequenced on an Illumina HiSeq 2500, using 50 bp SE reads (avg. number of reads = 30.5 million; see Supp. Table 3 for total reads per sample). We checked the quality of each file with *fastqc* (version 0.11.7) (20) and subsequently performed adapter- and quality-trimming with *cutadapt* (version 1.16; quality threshold of 20 and minimum retained length of 30 bp) (21). We used *STAR* (version 020201; (22)) to align RNAseq reads to each of the reference genomes on a per-exon basis, allowing for 50 bp of overhanging sequence on either side of the putative splice junctions (--sjdbOverhang 50). We sorted, indexed, and filtered (MQ > 2) bam files with *samtools* (version 1.6; (23)). To count RNAseq reads for each exon, we used *HTseq* (version 0.10.0) (24) with the stranded option set to “no” and a minimum quality of 0 (since bams were pre-filtered). For each exon, we calculated normalized expression as the number of transcripts per million (TPM), which was chosen based on its ability to compare across libraries (25). We filtered out any exons less than 50bp in length as this will influence our ability to map reads to these exons with our 50 bp reads.

Expression counts are available in the supplemental material (Supp. Files 3-5) Scripts used to prepare or generate these materials are available at https://github.com/HirschLabUMN/Split_genes/Per_Transcript_Exon_Pipeline.

### Syntenic homology pipeline

Identifying syntenic homologs across each annotation was done in two steps, which included identifying large blocks of synteny between the genomes and comparing specific BLAST searches within those large blocks (Figure 1A). For the first step, we used *nucmer* (version 3.1) for each pairwise combination of genomes (26), requiring anchor matches to be unique in both reference and query (“--mum” flag) as well as a minimum cluster length (--c) of 1000 bp. We used default settings for the remaining options. We ran the *delta-filter* utility within the *Mummer* suite to identify the longest mutually consistent set of matches (-g flag) with a minimum alignment uniqueness of 75% (-u flag). Finally, we used the *show-coords* utility to convert the output into a table set of coordinates.

For the second step, we began by performing an all-by-all BLAST (blastn) using the databases described in the previous section and retaining only the matches with an E-value < 1e-4. If there were multiple matches between a given query and subject gene pair, we kept only the single best match based on E-value (length of matching bases was used in case of equivalent E-values). We then filtered matches based on whether the subject and query CDS were within the same *nucmer*-established syntenic regions (± 500kb on each side). Lastly, we searched for instances where proximal subject genes (within five gene models as determined by numeric suffix of gene IDs) matched the same query gene. From this, we classified each query gene as having: 1.) no corresponding gene in the alternative annotation, 2.) a single corresponding gene, or 3.) multiple corresponding genes. We then looked for overlap among the reciprocal BLASTs to confirm syntenic homologous relationships. In the case of a single gene corresponding to multiple genes in one direction, we required that the multiple genes correspond exclusively to the single gene from the other reference. From the one-to-multiple syntenic homologies, we isolated the potential split-gene misannotations by requiring that the “multiple” genes are: 1.) not annotated as overlapping, 2.) on the same strand, 3.) not a tandem duplication based on BLAST (i.e. have less than 10% overlapping BLAST coordinates calculated as a percentage of total length covered by BLAST hits; L1 / L2 < 0.1, see Fig 1b for definition of L1 and L2), and 4.) each expressed in our dataset.

### Split-gene classification

Our classification method is based on the expectation that expression across the split genes should be less consistent if the split (multiple) gene annotation is correct than if the merged (single) gene annotation is correct. This implies two requirements: 1.) a metric that distills expression differences across split genes and 2.) critical values from a null distribution that specify values too large or small to be expected by chance.

For each gene we first calculated normalized expression (TPM) for every sequenced tissue (i.e. library) by averaging across exons, filtering out any genes in which this average is less than 0.01 (see Supp. Fig 6 for distribution of TPM values). For each split-gene candidate, we then calculated the log_2_-fold change in expression across all split genes within the set. We take the absolute value of this log_2_-fold change to erase the dependence on what is arbitrarily chosen as numerator and denominator (if we do not take the absolute values, then the distribution is centered on 0, as expected if expression is equivalent across split-genes; Supp. Fig. 10). If more than two genes correspond to a single, merged gene, we then averaged across all possible fold-change values to arrive at a single number summarizing expression differences across the split genes within a single tissue. The final metric for the split-gene candidate set is an average (across tissues and biological replicates) of these absolute log_2_-fold changes, which we term M2f for “mean two-fold expression change across tissues.”

When calculating this value, we subset the data to include only the genotype corresponding to the annotation with the split, or multiple gene models, in order to provide the best representation of the expression patterns used to create the annotation. If there is differential expression or differential exon usage between genotypes, then utilizing expression data from divergent genotypes could generate a false signal for M2f.

Next, we developed a simulation framework to generate empirical null distributions. The first distribution that we simulated was used to identify split-gene candidates whose expression differences are greater than we would expect by chance. Here, we randomly selected 17,583 total genes across the three annotations (excluding the split/merge candidates) and “split” them in two at a random position. We chose only genes with at least 4 exons for splitting to avoid simulating an overabundance of single-exon genes, though this minimum exon criteria did not have a large effect on the resulting distributions (Supp. Fig 3). We then calculated the M2f value across artificially split pairs to produce a distribution. Candidates with high M2f values relative to this distribution indicate that these genes show larger differences than we would expect if we were to simply take a truly merged gene and treat it as separate genes. We use the 90th percentile of the null distribution as the threshold to classify that split-gene candidates should in fact stay as separate genes.

The second null distribution was created by randomly choosing 48,408 total adjacent pairs of genes across the three annotations and artificially merging them into a single gene. We calculate the M2f values for the original adjacent loci and used the 10th percentile of the distribution for all M2f values for the artificially merged distribution as the threshold below which we classified split-gene candidates as merged (i.e. the single, merged gene model is correct). These distributions were similar across annotations of the different genotypes (Supp. Fig 4), with consequently similar values for the 10th percentile (1.12, 1.08, and 1.13 for B73, PH207, and W22, respectively) and 90th percentile (2.66, 2.52, and 2.79 for B73, PH207, and W22, respectively) of the simulated merged and simulated split distributions, respectively. Thus, for each of these percentiles, we used a single value (1.11 for 10th percentile and 2.66 for 90th percentile) based on pooling the simulated data across the annotations.

Input files are available in Supp. File 6-8, output file is available in Supp. File 9, and code used to prepare output from the syntenic homology pipeline and classify split-gene candidates is available at https://github.com/HirschLabUMN/Split_genes/tree/master/scripts.

### B73 IsoSeq and AED analysis

The PacBio IsoSeq data for B73 was downloaded from the SRA (BioProject #: PRJNA10769; SRA Project ID: SRP067440; SRA Sample Numbers: SRR3147022 through SRR3147057) (13). These FASTQ files contain intact cDNA fragments, which result from running the raw reads through the IsoSeq processing pipeline. For each of the six tissues, there were six FASTQ files corresponding to non-overlapping size ranges of the cDNA fragments. We mapped each fastq to the B73 v4 reference genome assembly, using the splice-aware settings in *Minimap2* (“-ax splice -uf -C5”) (27). We then combined all BAM files and filtered for cDNAs with a mapping quality > 20. We calculated coverage of each gene model using BEDtools (28), requiring that the IsoSeq cDNAs covered 75% of the gene model (according to the longest transcript; Sup. File 10).

Annotation Edit Distance (AED) values for the B73 (version 4) annotation were made available by the Ware lab at: ftp://ftp.gramene.org/pub/gramene/Zea_mays/Jamboree_materials/. AED scores for the PH207 annotation are provided in Supp. File 11.

### Differential expression and exon usage analysis

To investigate the effect of split-gene misannotations on differential expression and differential exon usage, we utilized the programs DEseq2 (version 1.22.2) (29) and DEXseq (version 1.28.3) (30), respectively. For each of these analyses, we were interested in determining whether conflicting biological conclusions would be drawn for one or more of the split-genes as compared to the single, merged gene and whether such conflicts occur at a higher rate for misannotated split-genes. We subsetted the data to include only the genotype that is not involved in the split-gene candidate set to avoid artifacts due to reference mapping bias. For example, if we are investigating two genes from the W22 annotation that correspond to a single gene in B73, then we would use only expression data from PH207 (mapped to both W22 and B73). If we used expression data from W22 (again, mapped to W22 and B73) and observed conflicting DE inference (e.g. DE for one of the W22 genes, but no DE for the B73 gene), we would be unable to disentangle whether the conflict was due to the misannotation or reference bias.

Since we are only utilizing expression data from a single genotype, we are restricted to testing for differential expression (or exon usage) across tissues. For DESeq2, we summed expression counts (non-normalized) across exons, filtered genes with no expression, and tested for differential expression with default parameters. For DEXseq, we directly used the per-exon expression counts from HTseq, again with default parameters. Our exact implementation of each of these analyses can be found at https://github.com/HirschLabUMN/Split_genes/blob/master/analysis/SplitGenes.Rmd.

## Declarations

### Ethics approval and consent to participate

Not applicable

### Consent for publication

Not applicable

### Availability of data and materials

The raw expression data supporting the conclusions of this article is available in the NCBI SRA repository (BioProject number: PRJNA543878). The input files, including the expression counts and the real and simulated split-gene candidates, are included as supplemental files. Code is available at https://github.com/HirschLabUMN/Split_genes/. The script that generated the tables, figures, and numbers is available at https://github.com/HirschLabUMN/Split_genes/blob/master/analysis/SplitGenes.Rmd

### Author contributions

CNH, NMS, SEM conceived the experiment. PJM, JMM, CO, AB analyzed the data. PJM wrote the manuscript. All authors reviewed and approved the final manuscript.

## Acknowledgements

We thank the lab of Doreen Ware (particularly, Marcela Tello-Ruiz, Josh Stein, and Cristina Fernandez-Marco) at Cold Spring Harbor Laboratory for sharing information on the B73 version 4 annotation as well as for organizing the Maize Annotation Jamboree, a workshop which enables community involvement in improving the B73 annotation.

## Supplemental Files

Supp. File 1: One-to-one homologs identified in the syntenic homology pipeline. Filename: recip_one2one_500kb.txt.

Supp. File 2: One-to-many homologs identified in the syntenic homology pipeline. A filtered subset of these entries are the split-gene candidates analyzed in this study. Filename: recip_one2many_500kb.txt

Supp. File 3: Expression matrix for reads mapped to B73. Sample names have format Genotype-Tissue-Rep. E.g. B-A-R1 is first biological replicate (R1) of anther tissue (A) from B73 (B). Filename: B73_ExpressionMatrix.txt

Supp. File 4: Expression matrix for reads mapped to W22. Sample names have format Genotype-Tissue-Rep. E.g. W-A-R1 is first biological replicate (R1) of anther tissue (A) from W22 (W). Filename: W22_ExpressionMatrix.txt

Supp. File 5: Expression matrix for reads mapped to PH207. Sample names have format Genotype-Tissue-Rep. E.g. P-A-R1 is first biological replicate (R1) of anther tissue (A) from PH207 (P). Filename: PH207_ExpressionMatrix.txt

Supp. File 6: Formatted input containing B73 candidate and simulated split-genes for classification via the M2f_Classify.R script. Filename: B73_split_genes.txt. Accompanying expression data is B73_ExpressionMatrix.txt

Supp. File 7: Formatted input containing W22 candidate and simulated split-genes for classification via the M2f_Classify.R script. Filename: W22_split_genes.txt. Accompanying expression data is W22_ExpressionMatrix.txt

Supp. File 8: Formatted input containing PH207 candidate and simulated split-genes for classification via the M2f_Classify.R script. Filename: PH207_split_genes.txt. Accompanying expression data is PH207_ExpressionMatrix.txt

Supp. File 9: Supported annotations according to our M2f procedure. Filename: M2f_Calls.txt

Supp. File 10: IsoSeq cDNA count data. See B73 IsoSeq analysis in Methods for procedure used to generate counts. Filename: IsoSeq.txt

Supp. File 11: Annotation Edit Distance (AED) scores for B73 and PH207 annotated gene models.

## Supplemental Tables and Figures

Supp. Table 1. Summary of overlapping annotations for candidate split-genes. The average overlap (Avg. Ovlp) is calculated as L1 / L2 in Fig 1b, except that in this case we are looking at the boundaries of the genes as they were originally annotated instead of the boundaries found when aligning these genes to an alternative reference genome.

Supp Figure 1. Features of split-gene candidates alongside one-to-one homologous genes. The merged candidates are the single, merged genes to which a set of split-gene corresponds. Exon density is calculated as the number of exons divided by the gene length. We required transcripts per million (TPM) > 0.01 for a gene to be considered expressed.

Supp Fig 2. M2f distributions for simulated merged genes, simulated split genes, and true split-gene candidates for multiple and single isoform genes in the B73 reference genome.

Supp. Fig 3. Distribution of M2f values for simulated split genes using different values for the minimum number of exons. B= B73, P = PH207, W= W22. An effect of exon number is seen on the dispersion of these distributions, but only for genes with very large exons (>20). Four exons is the median value for all annotations, which has a very similar distribution

Supp Figure 4. Null distributions from the simulated merged and simulated split genes for the three annotations. Differences in the distributions capture the cumulative effects of the annotation on our M2f metric. The 10th percentile of the simulated merge distribution was 1.16, 1.14, and 1.16 for B73, PH207, and W22, respectively. The 90th percentile for the simulated split distribution was 2.78, 2.66, and 2.97. Thus, the null distributions are relatively insensitive to the annotation used.

Supp. Figure 5. Additional features for comparison between one-to-one genes and split-gene candidates. Split-gene candidates are classified based on whether they were initially annotated as split or merged for a given genotype followed by the classification based on the M2f method. E.g. The “SS” box for the B73 Genotype are instances where multiple genes in B73 correspond to a single gene in either PH207 or W22 and the multiple (split) genes of B73 were determined to be the supported annotation. A.) Number of exons the gene is composed of. B.) Number of tissues with normalized expression > 0.01. C.) Exon Density is calculated as the number of exons divided by the gene length. D.) Proportion of genes in each category that are expressed (TPM < 0.01) in less than five tissues. E.) The key for the colors used in the figures (as in Fig 4A).

Sup Figure 6. Distributions of normalized expression for one-to-one genes as well as the split-gene candidates, which are further classified as to whether the original annotation was supported or not. (1-1 one-to-one gene; SS split is supported; SNS split is not supported; MS merged is supported; MNS merged is not supported).

Supp. Figure 7. Comparison of expression estimates across homologs. Incorrect split-gene annotations (split is not supported (SNS) instead of split is supported (SS), according to Figure 4). For the incorrect split-gene annotations (SNS), expression of each split-gene is compared to the expression value from the single gene model to which they correspond. Since the two split-genes actually correspond to the same underlying gene, we expect a strong correlation in expression between these genes.

Supp Figure 8. Distribution of adjusted p-values for differential expression across tissues for the single, merged genes from each annotation that correspond to multiple split-genes in an alternative annotation. MNS = Merged Not Supported (i.e. the multiple gene, split gene annotation is supported). MS = Merged Supported (i.e. the single, merged gene annotation is supported). There is a slight inflation of small p-values when the non-supported gene models are used.

Supp Figure 9. Distribution of the proportion of significant exons in a gene for DEXseq analysis. Differential exon usage is tested across tissues for the single, merged genes from each annotation that correspond to multiple split-genes in an alternative annotation. A p-value is reported per exon, and we required this to be < 0.05 to be counted as significant. MNS = Merged Not Supported (i.e. the multiple gene, split gene annotation is supported). MS = Merged Supported (i.e. the single, merged gene annotation is supported).

Supp Figure 10. M2f distributions when NOT taking the absolute value. Expected values of log2-fold changes should be centered on 0, indicating no difference in expression between split-genes.

