## Supplemental Tables and Figures for "Using multiple reference genomes to identify and resolve annotation inconsistencies"

Supp. Table 1. Summary of NUCMer for all pairwise combinations.  The Total Length is the summed length of all aligned sequences, where *n* is the total number of aligned sequences.

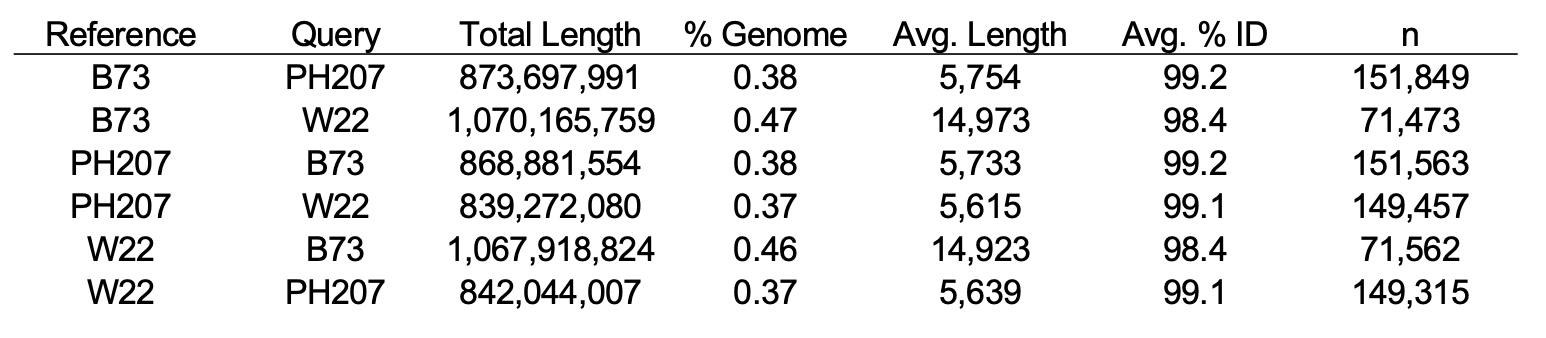

Supp. Table 2. Summary of overlapping annotations for candidate split-genes.  The average overlap (Avg. Ovlp) is calculated as L1 / L2 in Fig 1b, except that in this case we are looking at the boundaries of the genes as they were originally annotated instead of the boundaries found when aligning these genes to an alternative reference genome.

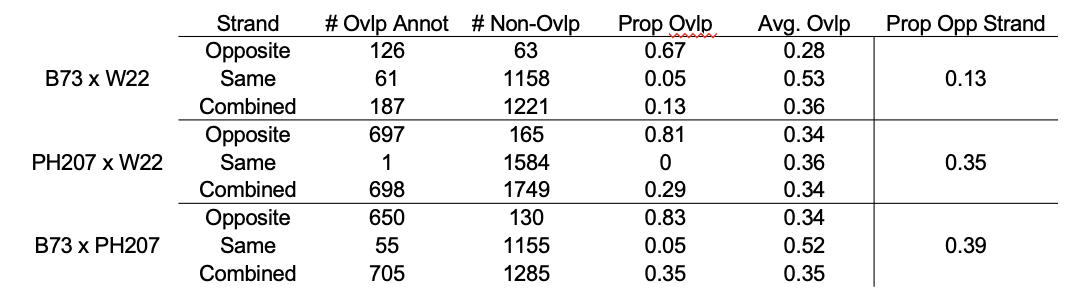

Supp. Table 3. Total read counts for each sample

| Sample | TotalReads |  | Sample | TotalReads |
| --- | --- | --- | --- | --- |
| B-A-R1 | 30612434 |  | P-L-R1 | 27470587 |
| B-A-R2 | 32891418 |  | P-L-R2 | 30177725 |
| B-Em-R1 | 26614422 |  | P-L10-R1 | 34909408 |
| B-Em-R2 | 29467766 |  | P-L10-R2 | 34024762 |
| B-En-R1 | 23156229 |  | P-R-R1 | 28559074 |
| B-En-R2 | 32261390 |  | P-R-R2 | 40864817 |
| B-I-R1 | 28752577 |  | P-SC-R1 | 30026306 |
| B-I-R2 | 31380132 |  | P-SC-R2 | 31428071 |
| B-IE-R1 | 27464746 |  | P-T-R1 | 31588449 |
| B-IE-R2 | 27898833 |  | P-T-R2 | 32123964 |
| B-L-R1 | 26810281 |  | W-A-R1 | 25084628 |
| B-L-R2 | 27453240 |  | W-A-R2 | 28264792 |
| B-L10-R1 | 34585599 |  | W-Em-R1 | 29422112 |
| B-L10-R2 | 28978662 |  | W-Em-R2 | 27085556 |
| B-R-R1 | 30219442 |  | W-En-R1 | 28349977 |
| B-R-R2 | 31281356 |  | W-En-R2 | 35811617 |
| B-SC-R1 | 31399874 |  | W-I-R1 | 28641145 |
| B-SC-R2 | 30194385 |  | W-I-R2 | 32153246 |
| B-T-R1 | 30650037 |  | W-IE-R1 | 25521054 |
| B-T-R2 | 33406864 |  | W-IE-R2 | 33543447 |
| P-A-R1 | 31084506 |  | W-L-R1 | 27623687 |
| P-A-R2 | 27760591 |  | W-L-R2 | 30749796 |
| P-Em-R1 | 31523525 |  | W-L10-R1 | 31710277 |
| P-Em-R2 | 32847687 |  | W-L10-R2 | 30738420 |
| P-En-R1 | 31455071 |  | W-R-R1 | 32573086 |
| P-En-R2 | 35522111 |  | W-R-R2 | 33121493 |
| P-I-R1 | 28779102 |  | W-SC-R1 | 29014567 |
| P-I-R2 | 31767169 |  | W-SC-R2 | 28749269 |
| P-IE-R1 | 30539071 |  | W-T-R1 | 30501912 |
| P-IE-R2 | 32379450 |  | W-T-R2 | 30582209 |

Supp Figure 1.  Features of the split-gene candidates alongside the One-to-one homologous genes.  The Merged Candidates are the single, merged genes to which a set of split-gene corresponds to.  Exon Density is calculated as the number of exons divided by the gene length. We required TPM > 0.01 for a gene to be considered expressed.

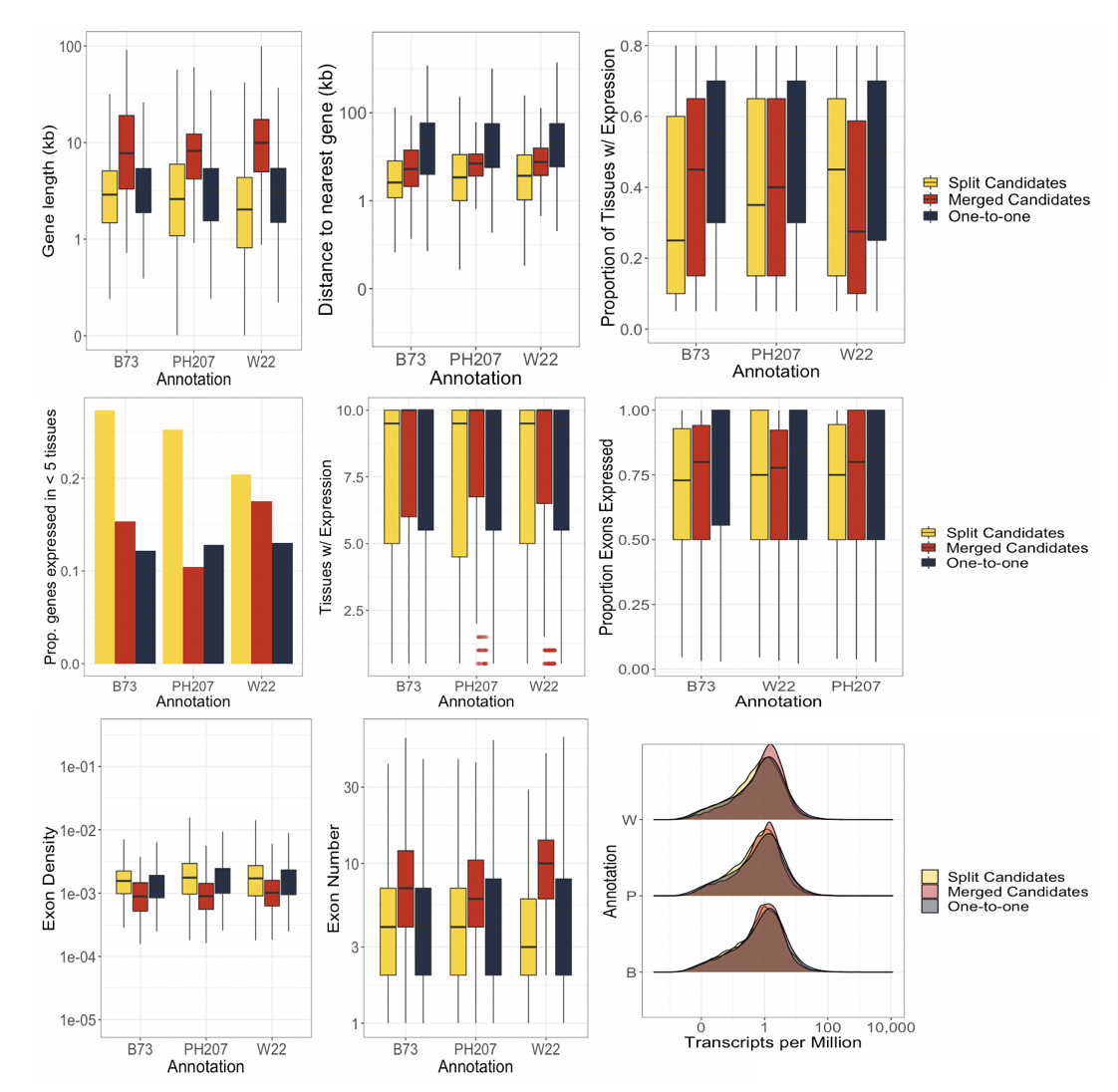

Supp. Fig 2. M2f distributions for simulated merged genes, simulated split genes, and true split-gene candidates for multiple and single isoform genes in the B73 reference genome.

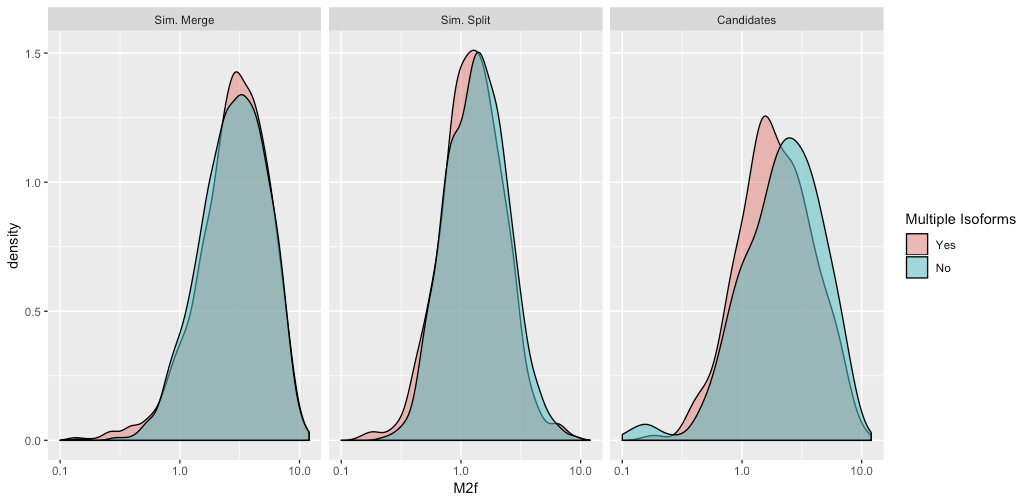

Supp. Fig 3. Distribution of M2f values for simulated split genes using different values for the minimum number of exons. B= B73, P = PH207, W= W22.  An effect of exon number is seen on the dispersion of these distributions, but only for genes with very large exons (>20). 4 exons is the median value for all annotations, which has a very similar distribution

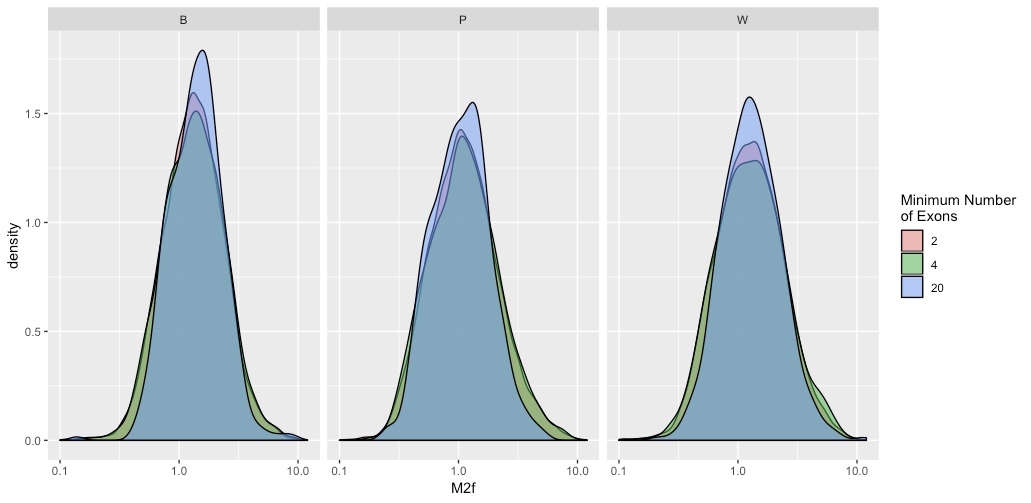

Supp Figure 4. Null distributions from the simulated merged and simulated split genes for the 3 annotations.  Differences in the distributions capture the cumulative effects of the annotation on our M2f metric. The 10th percentile of the former was 1.16, 1.14, and 1.16 for B73, PH207, and W22, respectively.  The 90th percentile for the latter was 2.78, 2.66, and 2.97. Thus, the null distributions are relatively insensitive to the annotation used.

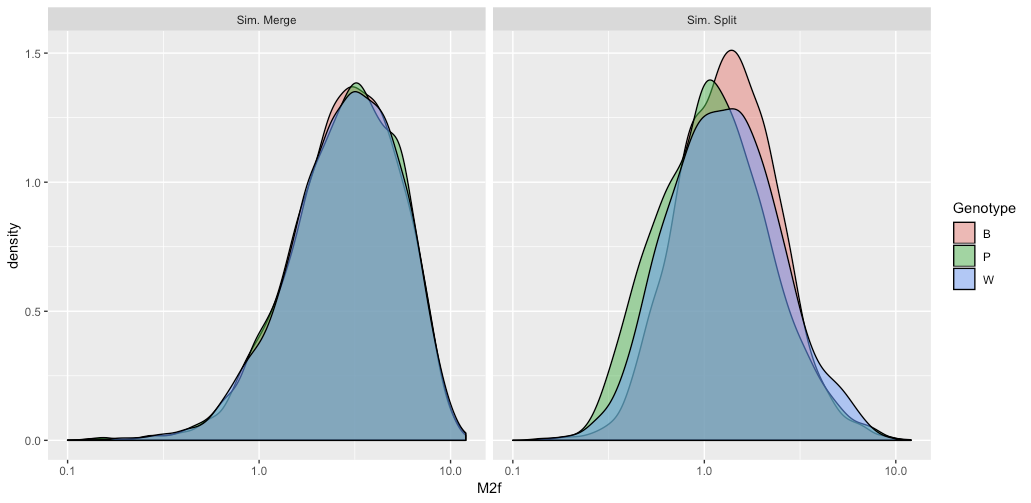

Supp. Figure 5.  Additional features for comparison between One-to-one genes and split-gene candidates.  The latter are classified based on whether they were initially annotated as split or merged for a given genotype followed by the classification based on the M2f method.  E.g. The “SS” box for the B73 Genotype are instances where multiple genes in B73 correspond to a single gene in either PH207 or W22 and the multiple (split) genes of B73 were determined to be the supported annotation.  A.) Number of exons the gene is composed of. B.) Number of tissues with normalized expression > 0.01. C.) Exon Density is calculated as the number of exons divided by the gene length. D.) Proportion of genes in each category that are expressed (TPM < 0.01) in less than five tissues.  E.) The key for the colors used in the figures (as in Fig 4A).

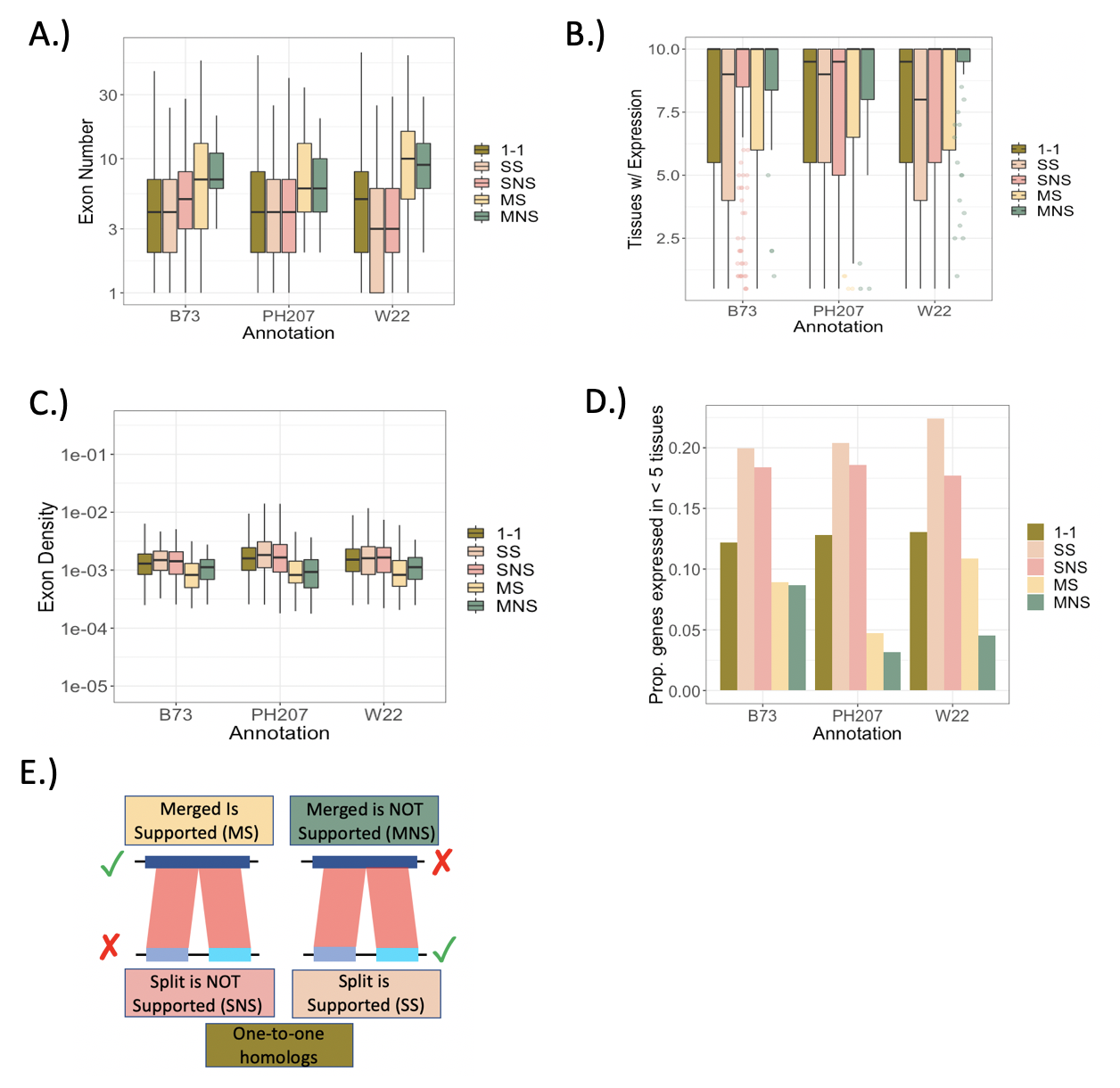

Supp. Figure 6.  Distributions of normalized expression for one-to-one genes as well as the split-gene candidates, which are further classified as to whether the original annotation was supported or not.

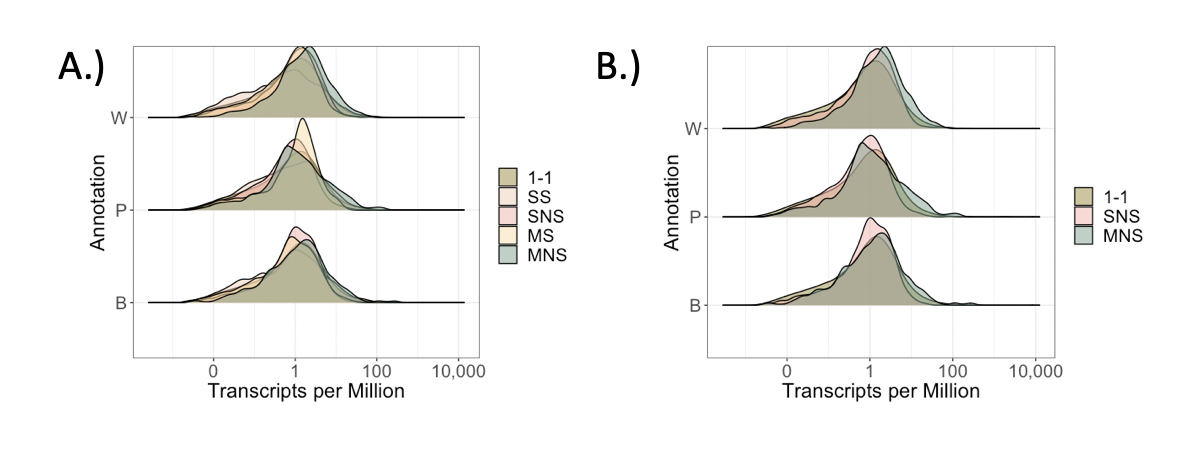

Supp. Figure 7.  Comparing expression estimates across homologs, similar to Figure 5A. Here, we plot the ***in***correct split-gene annotations (SNS instead of SS, according to Figure 4).  For the ***in***correct split-gene annotations, expression of each split-gene is compared to the one expression value from the single gene that they correspond to.  Since the two split-genes actually correspond to the same underlying gene, we expect a strong correlation in expression between these genes.

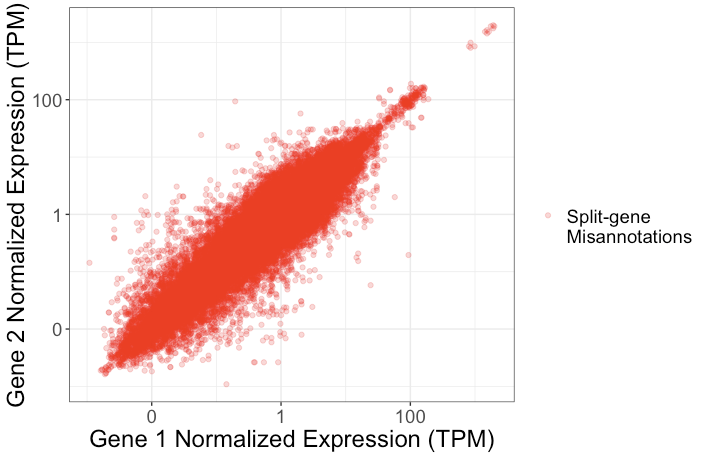

Supp. Figure 8.  Distribution of adjusted p-values for differential expression across tissues for the single, merged genes from each annotation that correspond to multiple split-genes in an alternative annotation.  MNS = Merged Not Supported (i.e. the correct annotation is for multiple genes as in the alternative annotation). MS = Merged Supported (i.e. the single, merged gene annotation is supported). There is a slight inflation of small p-values when the non-supported gene models are used.

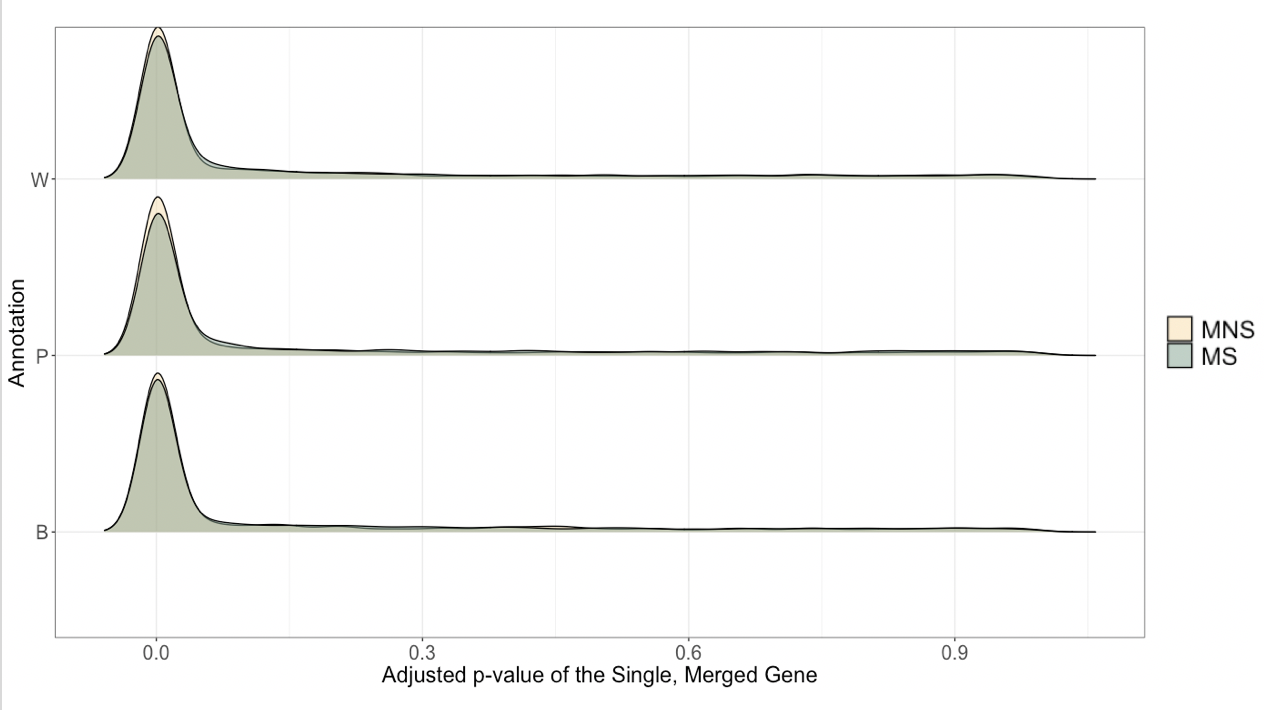

Supp. Figure 9.  Distribution of the proportion of significant exons in a gene for DEXseq analysis.  Differential exon usage is tested across tissues for the single, merged genes from each annotation that correspond to multiple split-genes in an alternative annotation.  A p-value is reported per exon, and we required this to be < 0.05 to be counted as significant. MNS = Merged Not Supported (i.e. the supported annotation is for multiple genes as in the alternative annotation).  MS = Merged Supported (i.e. the single, merged gene annotation is supported).

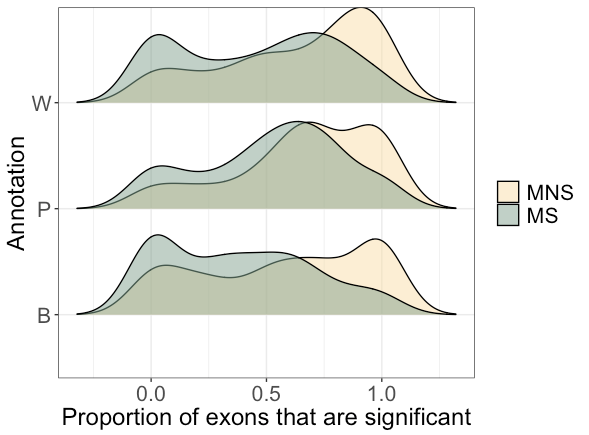

Supp. Figure 10.  M2f distributions when NOT taking the absolute value.  Expected values of log2-fold changes should be centered on 0, indicating no difference in expression between split-genes.

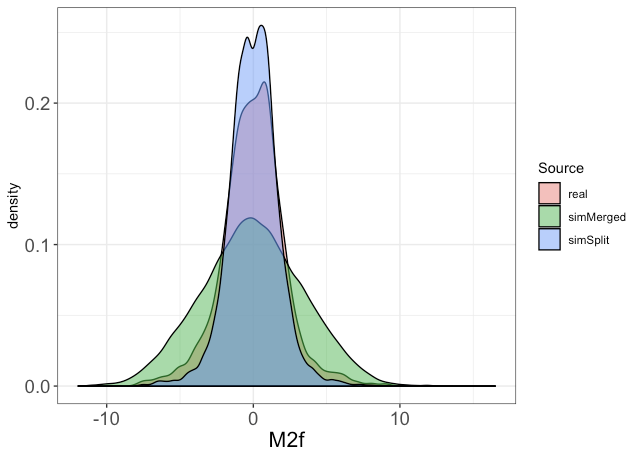
